# Respiration modulates oscillatory neural network activity at rest

**DOI:** 10.1101/2020.04.23.057216

**Authors:** Daniel S. Kluger, Joachim Gross

## Abstract

Despite recent advances in understanding how respiration affects neural signalling to influence perception, cognition, and behaviour, it is yet unclear to what extent breathing modulates brain oscillations at rest. We acquired respiration and resting state magnetoencephalography (MEG) data from human participants to investigate if, where, and how respiration cyclically modulates oscillatory amplitudes (2 – 150 Hz). Using measures of phase-amplitude coupling, we show respiration-modulated brain oscillations (RMBOs) across all major frequency bands. Sources of these modulations spanned a widespread network of cortical and subcortical brain areas with distinct spectro-temporal modulation profiles. Globally, high-frequency gamma modulation increased with distance to the head centre, whereas delta and theta modulation decreased with height in the sagittal plane. Overall, we provide the first comprehensive mapping of RMBOs across the entire brain, highlighting respiration-brain coupling as a fundamental mechanism to shape neural processing within canonical resting-state and respiratory control networks.

## Introduction

We all breathe. Human respiration at rest comes naturally and comprises active (but automatic) inspiration and passive expiration ^1^. The rhythmicity of each breath is initiated and coordinated by coupled oscillators periodically driving respiration, most prominently the preBötzinger complex located in the medulla ^2^. This microcircuit typically controls respiration autonomously, making the act of breathing seem effortless. Importantly, however, respiration is also under top-down control, as evident from adaptive breathing during e.g. speaking, laughing, and crying ^3^. Hence, there is a bidirectional interplay of the cortex and rhythmic pattern generators of respiration: Efferent respiratory signals from the preBötzinger complex project to suprapontine nuclei (via locus coeruleus and olfactory nuclei ^4^) as well as to the central medial thalamus, which is directly connected to limbic and sensorimotor cortical areas ^5^. In turn, cortical areas evoke changes in the primary respiratory network, e.g. to initiate specific motor acts (e.g. swallowing or singing) or transitions between brain states (e.g. during panic attacks).

As neural oscillations have been established as sensitive markers of brain states in general ^6^, the question arises to what extent rhythmic brain activity is modulated by the rhythmic act of breathing. Indeed, studies of respiration-brain coupling have recently attracted increased attention, reporting a range of cognitive and motor processes to be influenced by respiration phase. Human participants were found to spontaneously inhale at onsets of cognitive tasks ^7^ and respiration phase modulated neural responses in sensory ^8^ and face processing ^9^ tasks as well as during oculomotor control ^10^ and isometric contraction ^11^. Parallel to this body of work, animal studies have conclusively shown respiration to entrain brain oscillations not only in olfactory regions ^12^, but also in rodent whisker barrel cortex ^13^ and even hippocampus ^14^. In other words, brain rhythms previously attributed to cognitive processes such as memory were demonstrated to at least in part reflect processes closely linked to respiration ^15^.

Despite significant advances in the animal literature, these links are still critically understudied in humans. Notable exceptions include intracranial EEG (iEEG) work in epileptic patients corroborating that oscillations at various frequencies can be locked to the respiration cycle even in non-olfactory brain regions ^9^. Moreover, two non-invasive studies recently linked respiration phase to changes in task-related oscillatory activity ^7^. Overall, both animal and human studies all lead to three fundamental questions that recognise respiration as a vital, continuous rhythm persisting during all tasks and activities as well as at rest: i) to what extent does breathing modulate rhythmic brain oscillations at rest, ii) where are these modulatory effects localised in the brain, and iii) how does modulation unfold over the course of the respiration cycle. Therefore, what is needed is a comprehensive account integrating recent findings of respiration-brain coupling against the anatomical backdrop of canonical resting state and respiratory control networks (RCN). A variety of neural networks have extensively been described to organise the brain’s intrinsic or ongoing activity, among which the default mode network (DMN), the dorsal attention network (DAN), and the salience network (SN) have received particular attention ^16^. Previous studies have demonstrated intriguing anticorrelated dynamics of activity between these large-scale networks (i.e., increases in one network lead to decreases in another ^17^). Such fluctuating relationships between cortical networks could conceivably be modulated by changes in body states such as respiration. The full picture is complemented by equally prominent networks of deeper sites known to be involved in respiratory control. In addition to pattern generators like the preBötzinger complex in the medulla, other subregions within the brain stem ^18^ as well as cerebellar nuclei ^19^ and, naturally, olfactory areas ^12^ have been shown to be associated with the act of breathing. Interestingly, the respiratory control network also includes directly connected cortical sites like primary and supplementary motor areas ^20^ and even shows anatomical overlap with resting state networks, namely within medial prefrontal cortex ^21^, insula ^22^, and ACC ^23^. We thus aimed to investigate respiration-related modulations of oscillatory brain activity and its spectro-temporal characteristics at rest, relating their anatomical sources to canonical networks of both resting state activity and respiratory control.

To this end, we simultaneously recorded spontaneous respiration and eyes-open resting state MEG data from healthy human participants. Using the modulation index (MI) as a measure of crossfrequency phase-amplitude coupling ^24^, we first assessed respiration-induced modulation of brain oscillations globally across the entire brain. We then extracted single-voxel time series to localise the anatomical sources of these global modulation effects using beamforming (Fig. 1a). We employed non-negative matrix factorisation (NMF) for dimensionality reduction, effectively yielding a spatially constrained network of cortical and subcortical sources of respiration phase-dependent changes in rhythmic brain activity. Finally, we identified distinct spectro-temporal profiles of network components, highlighting an intriguing organisational pattern behind respiration-induced modulation of neural oscillations across the brain.

**Fig. 1.**
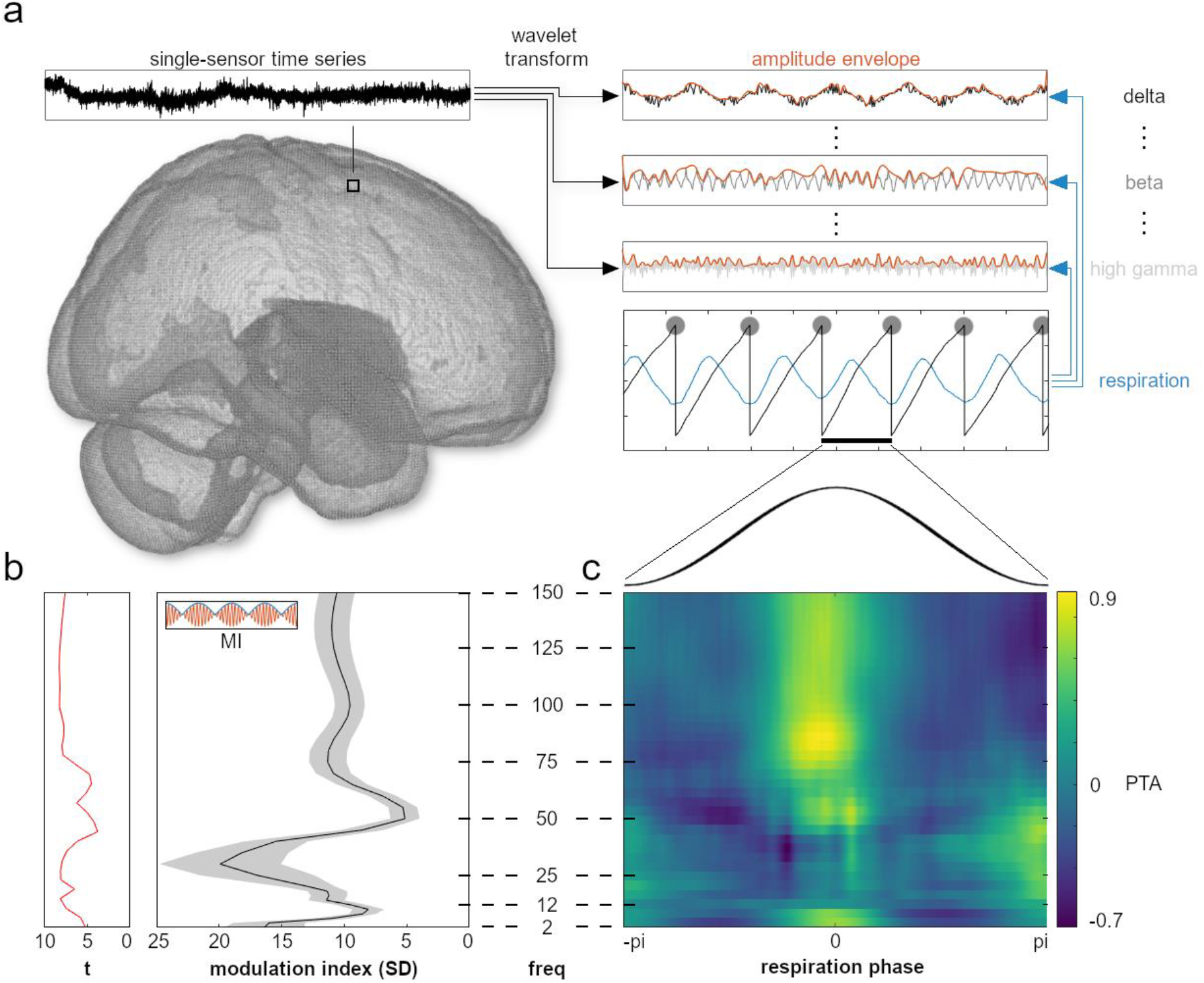
Respiration-induced modulation of global field power. **a**, Exemplary schematic of our analysis approach showing the wavelet transform of time series data from each voxel. Global field power was computed on the time course of all 268 channels for further analyses of the modulation index (MI) and phase-triggered average (PTA). MI quantifies to what extent the amplitude envelopes of frequency-specific brain oscillations (top right, red) were modulated by respiration (centre right, blue). This way, we computed modulation indices for each sensor, frequency, and participant before localising voxelwise time series in source space (see Fig. 2). **b**, Mean normalised MI (± SEM) over the entire frequency spectrum (right) and corresponding t-values from the cluster permutation test (left). Random shifts of respiration phase were employed to correct for low-frequency bias and to express MI in units of standard deviation (SD) of a surrogate distribution (leading to normalised MI, see Methods section). **c**, Mean PTAs across the respiratory cycle over the entire frequency spectrum. PTAs were computed by averaging frequency-specific amplitude envelopes (panel a) time-locked to peak inhalation. Note that PTAs were standardised for illustration purposes. Therefore, they show relative changes over the respiration cycle within each frequency band and do not directly correspond to absolute MI amplitudes from panel b.

## Results

### Respiration phase modulates global field power

To assess the fundamental question of whether respiration modulates oscillatory brain activity at rest, we first computed the modulation index (MI) in sensor-space (using all 268 channels) for whole-brain global field power ranging from 2 – 150 Hz. This analysis quantifies to what extent the amplitude of global brain oscillations is modulated by the phase of respiration. Our cluster permutation analysis revealed significant respiration-locked modulation of global field power indicated by the high normalised MI across the entire frequency spectrum (all *p* < .001, FDR-corrected at α = .05; see Fig. 1b). Local peaks with strongest modulation occurred at about 2, 30, 75, and 130 Hz (with strongest absolute modulation effects in the beta band), indicating differential modulation of specific brain oscillations. Next, we computed the phase-triggered average (PTA) to characterise these global modulation effects over a respiratory cycle. PTA is computed as the average of oscillatory amplitude across windows centered on all time points of peak inhalation. We found respiration phase to differentially modulate oscillations of various frequencies with distinct time courses. In particular, whereas most frequency bands showed strongest modulation effects around the inspiration peak, beta oscillations were visibly coupled to a different phase of respiration around inspiration onset (Fig. 1c).

This first analysis therefore revealed that the amplitude of global oscillatory brain activity was significantly modulated by respiration in a broad frequency range from 2 to 150 Hz with a temporal modulation that differs across frequencies. To gain a deeper understanding of how respiration modulates rhythmic activity across the brain, two questions immediately ensued, namely to localise the anatomical sources of such modulation effects and to explore their spectro-temporal profiles in more detail.

### Modulatory effects of respiration phase can be traced to cortical and subcortical networks

To identify the anatomical sources of these global modulations, we quantified how strongly respiration modulated the amplitude of brain oscillations within each voxel in the brain of each participant at each frequency between 2 and 150 Hz by computing the modulation index (MI). Next, we used sparse non-negative matrix factorisation (NMF) to reduce the dimensionality of the three-dimensional data set (participants x voxels x frequency; see Methods section). This resulted in an optimal low-dimensional representation consisting of 17 components. Each component reflected respiration-modulated brain oscillations (RMBOs) across the frequency spectrum within a specific brain area (either cortical or subcortical). For all 17 components, we show the spatial location of the network on an inflated brain, PTAs that illustrate the modulation of its oscillatory activity across the respiration cycle, and the full MI spectrum with shading corresponding to frequency bands of significant modulation (all *p* < .002, FDR-corrected across frequencies and components at α = .05; see Fig. 2). Together, this provides a comprehensive spatio-temporo-spectral account of respiration-modulated networks in the resting brain.

**Fig. 2.**
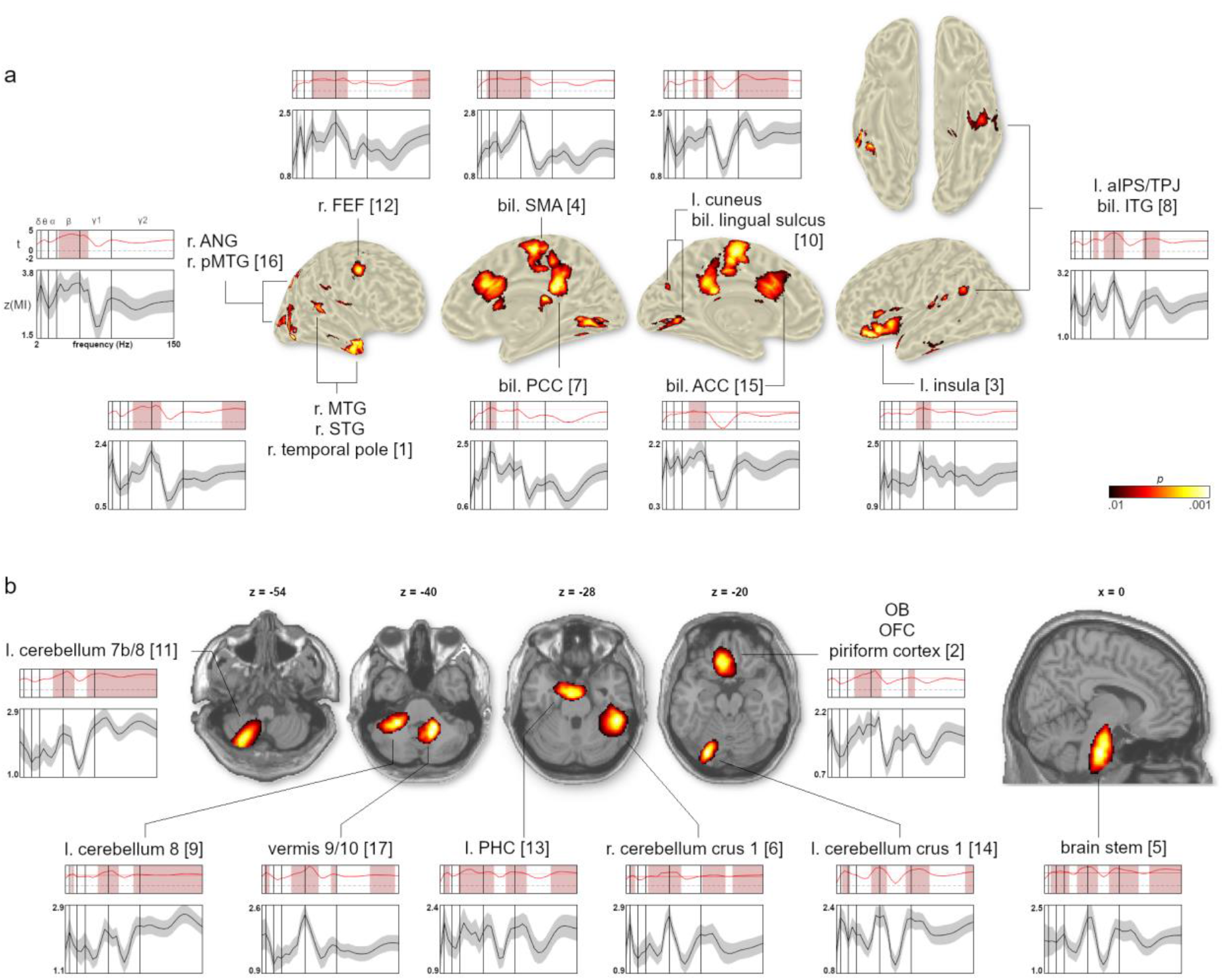
Anatomical locations and spectral modulation profiles of NMF components whose neural oscillations were significantly modulated by respiration. **a**, Cortical components plotted on an inflated brain surface. Bottom graphs illustrate each component’s normalized average MI course across frequencies (2 – 150 Hz). Upper graphs show component-specific t-value spectra from the cluster permutation test (with significant FDR-corrected frequencies shaded). Horizontal red line marks the significance threshold of each component, vertical lines mark borders between frequency bands (delta to high gamma). **b**, Subcortical components plotted on transverse and sagittal slices of the MNI brain. Same format as **a**.

Figure 2a shows the network’s cortical sources to be localised along the midline (ACC, SMA, PCC, cuneus, lingual sulcus) as well as in lateralised frontal (FEF, insula), temporal, and parietal cortices (angular gyrus, IPS). The network’s deeper, subcortical sites included several lateralised (crus 1, lobules 7b/8) and midline (vermis 9/10) subsections within the cerebellum, left parahippocampal cortex, and medial sources in the olfactory bulb/OFC (extending into piriform cortex) and brain stem (Fig. 2b).

These results provide several important insights. First, respiration significantly modulates oscillatory brain activity within a specific, but widely distributed cortical and subcortical brain network. Second, across these areas significant RMBOs can be found across almost the entire frequency range from 2-150 Hz. Third, time-frequency PTA maps demonstrate that the temporal modulation pattern of RMBOs is by no means uniform across frequencies and brain areas. Instead, some RMBOs are strongest at peak inhalation with others peaking at maximum expiration or at different times in between.

### Distinct spectro-temporal profiles of RMBO sites

Having localised the anatomical network underlying RMBOs, we next attempted to map distinct modulation patterns to anatomical subnetworks, with similarly modulated sites being grouped together. To this end, we employed hierarchical clustering of all 17 network components based on their modulation index across the frequency spectrum (as shown in Fig. 2). This data-driven approach yielded a total of seven clusters comprising between one and five components (Fig. 3, see Supplementary Fig. 4 and Methods for details): Cluster A consisted of a single component within the left insular cortex and showed significant modulation in the beta band. Cluster B showed a clear cortical organisation along the midline, comprising two bilateral PCC and SMA components with significant modulation from theta to low gamma oscillations. Similarly, cluster C comprised two components within bilateral ACC and FEF with significant modulation from delta to alpha and low gamma oscillations. Cluster D was formed by a total of five components spanning temporal (STG, MTG, ITG) and parietal cortices (aIPS/TPJ, angular gyrus) as well as deep cerebellar areas showing RMBOs. Due to its widespread topography, at least one cluster component showed significant modulation across the entire frequency spectrum. Cluster E again consisted of a single component (spanning left cuneus und bilateral lingual sulcus) with significant modulation in the delta and theta as well as in the low and high gamma band. Finally, two clusters were formed exclusively by deep sources: Cluster F comprised two components within the left cerebellum where low and high gamma oscillations were significantly modulated by respiration. Cluster G consisted of four components within the left parahippocampal cortex, brain stem, cerebellum, and olfactory bulb/OFC and showed significant modulation in the delta, theta, beta, and low gamma band. For more details on clustering results, see Supplementary Fig. 4.

**Fig. 3.**
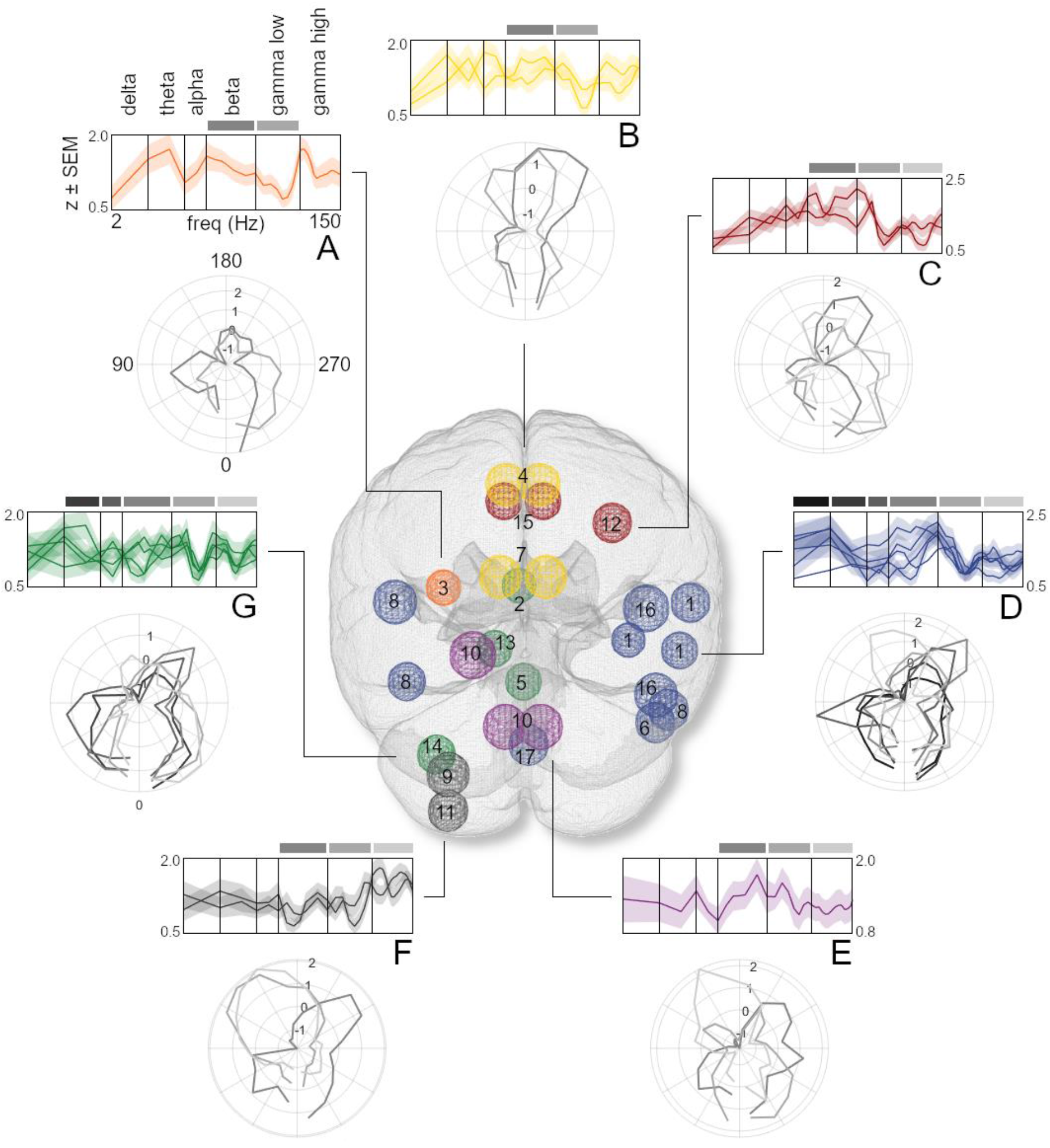
Time-frequency characteristics and anatomical distribution of component clusters. Hind view of the glass brain illustrates the spatial distribution of component clusters A-F (numeration according to Fig. 2; see Supplementary Fig. 5 for top and side views). Spheres mark peak locations of components and are colored according to cluster affiliation. Top curve plots depict z-transformed modulation indices of individual components within the cluster (group-level mean ± SEM) over the log-transformed frequencies. Vertical bars mark borders between frequency bands, superimposed horizontal bars indicate frequency bands in which at least one of the cluster’s components showed significant modulation due to respiration phase (also see Fig. 2). Polar plots illustrate the temporal modulation of RMBOs of these frequency bands averaged within clusters as a function of respiration phase (shown in angular units where 180° corresponds to the peak of the respiration signal). Significant frequencies are shown in a grey scale that codes frequency bands (from black (delta band) to light grey (high gamma band)).

The components’ spectral profiles revealed two noticeable links between anatomic location and oscillatory modulations: First, lateral components appeared to show stronger modulation of high frequencies (particularly within the high gamma band) than those closer to the head centre. Second, low frequencies (particularly within the delta band) appeared to be more strongly modulated within components located low in the sagittal plane compared to those located higher on the z axis. Linear mixed effect models corroborated these relationships, showing that the fixed effect of distance to the head centre significantly influenced high gamma (*t*(474) = 3.55, *p* < .001) but not low gamma modulations (*t*(474) = 1.57, *p* = .12) with stronger modulations for more lateral components. Moreover, height in the sagittal plane was found to significantly reduce modulatory effects on delta (*t*(474) = −5.21, *p* < .001) and theta oscillations (*t*(474) = −3.25, *p* = .001), demonstrating stronger low-frequency modulation for deeper brain areas.

Intriguingly, not only were different frequency bands modulated within a network of cortical and subcortical sites, but the time courses of these modulatory effects were equally frequency-specific. Polar plots in Figure 3 show the temporal modulation of RMBOs across the respiratory cycle for each cluster. Respiration phase was differentially coupled with amplitudes of low-frequency oscillations (such as delta and theta) compared to high-frequency oscillations (e.g. within the gamma band). Low frequencies consistently showed higher amplitudes during the beginning and end of a respiration cycle (with lowest amplitudes around the respiration peak), whereas the pattern appeared reversed for higher frequencies (see Figs. 3 and 4d as well as Supplementary Fig. 6). While specific spatio-temporal interactions of respiration-brain coupling exceeded the conceptual scope of this study, our findings are the first to suggest such spatio-spectral gradients and thus warrant detailed examination in future work.

**Fig. 4.**
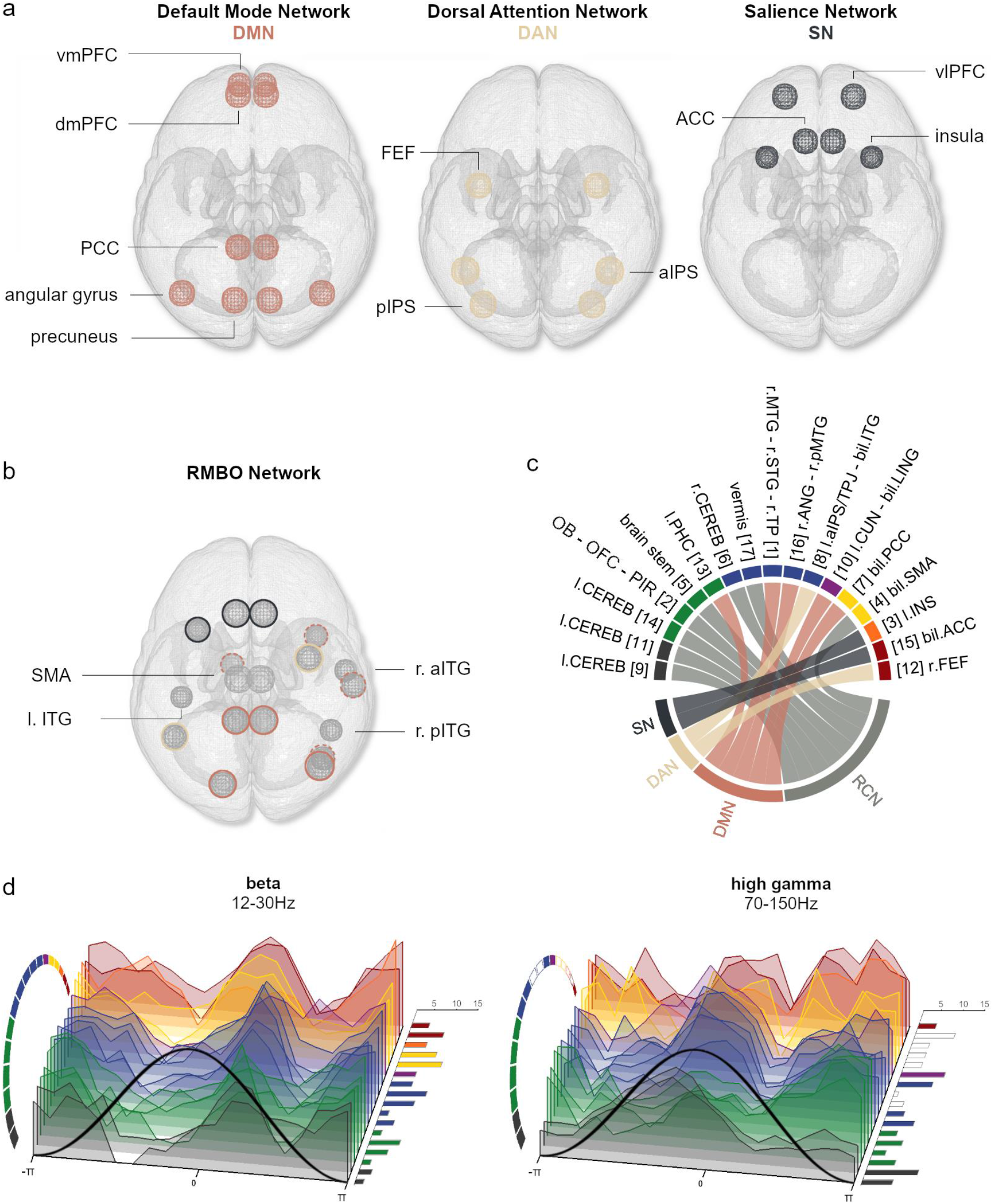
Mapping clusters of NMF components to canonical neural networks. **a**, Top-view stylised illustrations of neural nodes composing the default mode network (DMN), dorsal attention network (DAN), and salience network (SN) as described in the literature. **b**, Cortical brain areas showing significant RMBOs (as in Fig. 3) are colour-coded according to their correspondence to the resting state networks shown in **a**. As the MTG has increasingly been included in the DMN but was not part of its original formulation, NMF components located within the MTG are marked with a dashed line. Components within SMA and ITG were the only cortical sites not mapping to resting state networks (but see the Discussion for SMA as an established site of respiratory control). **c**, Direct mapping of all 16 clustered NMF components to the resting state neural networks (see **a**) and the respiratory control network (RCN) gained from the literature. Colour code for clusters A – G taken from Fig. 3. **d**, Waterfall plots show z-transformed amplitude modulation phase-locked to the respiration cycle exemplified for beta (left) and high gamma oscillations (right; see Supplementary Figure 6 for remaining frequency bands). Clusters of NMF components are shown in the same order as in **c**. Right-panel bar graphs show the number of participants whose modulation within the respective component was strongest for the depicted frequency band (vs all other frequency bands). Coloured bars and circular segments mark NMF components for which the respective frequency band was significantly modulated by respiration phase.

### RMBOs within nodes of resting state and respiratory control networks

Extending the distinction of deep vs more superficial components, cortical RMBOs were predominantly found in brain areas that make up canonical resting state networks (shown in Fig. 4a). With the exception of components within SMA (which has been shown to control respiratory functioning, see below) and ITG, all cortical modulation sites have previously been established as nodes within the DMN (PCC, angular gyrus, precuneus), DAN (FEF, aIPS), or SN (insula, ACC; see ^15^). Moreover, all deep and cerebellar modulation sites corresponded to a mostly subcortical network of brain areas controlling respiratory function, including bilateral cerebellum, olfactory bulb/OFC, brain stem, and SMA (Fig. 4c). Finally, Figure 4d suggests that although RMBOs of different frequencies had distinct temporal modulation profiles in general, there could also be certain sequential modulation patterns across clusters within a particular frequency. For example, while significant modulation of beta oscillations showed a general peak around expiration onset (distinct from e.g. high gamma modulation), this peak appeared to occur earlier and less pronounced in cluster B (PCC, SMA) than in cluster D (vermis, MTG, ANG). Overall, our results provide a unique perspective on the link between respiration phase and changes in oscillatory activity, mapping the sources of these modulatory effects to nodes of canonical networks in control of resting state activity and respiratory function.

## Discussion

Using non-invasive MEG recordings of human participants at rest, we performed the first spatially and spectrally comprehensive analysis of brain activity that is modulated by respiration. We identified respiration-modulated brain oscillations (RMBOs) across the entire spectrum between 2 and 150 Hz within a widespread network of cortical and subcortical brain areas. Intriguingly, instead of a uniform modulation pattern across brain areas and frequencies, our analysis revealed respiratory modulation signatures that differed between brain areas in frequency and the temporal modulation profile. Our results demonstrate that respiration significantly modulates oscillatory brain activity in a manner that is precisely orchestrated across frequency bands and networks of resting state activity and respiratory control. In what follows, we will integrate our novel results with the existing animal and human literature, characterise the functionality of neural oscillations within distinct networks, and attempt to provide an overview of potential multi-level mechanisms behind RMBOs.

### Subcortical and cortical sites of respiration-brain coupling

Gamma oscillations within the olfactory bulb (OB) were the first to be described in detail ^25^ and reflect local computations within OB ^26^. In a next step, slower (e.g. beta band) oscillations are thought to organise such local activity across brain areas ^27^ and appear to be the most coherent within OB ^28^. Similarly, even slower theta oscillations play a crucial role in the temporal organisation of neural activity within the hippocampal network and, consequently, its coordination with the mPFC ^29^. Our findings substantially advance these notions by showing that respiration phase modulates both low and high oscillatory frequencies within a spatial array comprising OB/OFC, brain stem, and cerebellum. As described earlier, the preBötzinger complex is widely regarded as the main pattern generator of respiratory rhythms ^4^ within the brain stem and several studies have implicated surrounding medullar as well as cerebellar sites in mediating voluntary control of respiration ^3^. This functional co-activation stems from ascending catecholaminergic neurons in the brain stem receiving projections from the cerebellum, particularly the vermis ^30^. In addition to brain stem projections, the vermis regulates autonomic responses including cardiovascular tone and respiration through connections to the spinal cord and hypothalamus ^31^. Going back to gamma oscillations, the cerebellum has been suggested to use cerebello-thalamic pathways to exert its effect on cortical gamma activity, especially within sensorimotor cortices ^32^: The cerebellum sends projections to the ‘motor’ ventral anterior lateral nucleus (VAL) of the thalamus, the (higher-order) posterior thalamic nucleus (VP, connected to primary motor and sensory cortices), and intralaminar nuclei, allowing cerebello-thalamic connections to coordinate and synchronise gamma oscillations across cortical areas ^33^.

Well-established bidirectional connections exist between the cerebellum and the neocortex via medullar and thalamic pathways. The cerebellum itself projects to motor and nonmotor cortical areas, including prefrontal and posterior parietal cortex ^34^. In turn, it receives input from a wide range of higher-order, nonmotor areas within the extrastriate cortex, posterior parietal cortex, cingulate cortex, and the parahippocampal gyrus ^34^, which is monosynaptically connected to the OB ^35^. Linking cortical and deep nodes of the RMBO network, distinct thalamic pathways can be found within the thalamus: The anterior thalamus is strongly connected to the SMA, premotor and (pre-) frontal cortex, and the ACC. A similar connectivity profile has been shown for the insular cortex ^18^. In sum, our findings integrate and extend a variety of individual results in two ways: First, cortical nodes within the RMBO network precisely reflect bidirectional projection areas of the deep and subcortical nodes (OB, brain stem, and cerebellum) via medullar and thalamic pathways. Second, the cortical nodes markedly resemble ‘sensorimotor distributions’ shown in multiple fMRI studies of respiratory control ^36^, raising the question as to how different cortical areas – motor areas, ACC, and insular cortex – are involved in the act of breathing. As both premotor and supplementary motor cortices contain representations of respiratory muscles ^37^, they have long been implicated in respiratory control. Similarly, ACC has been identified in studies of air hunger ^23^ and CO_2_-stimulated breathing. Finally, insular cortex activation is a consistent feature of many neuroimaging studies of dyspnoea ^22^. The close mapping of frontal, cingulate, and parietal areas to canonical resting state networks (see Fig. 4) suggests a general involvement of respiration in human brain processing irrespective of particular task demands. In this context it is noteworthy that nodes of resting state networks exhibit amplitude correlations predominantly in the beta frequency band ^38^. In our data, this frequency band shows strongest global modulation by respiration (Fig. 1b) and features prominently in the coupling of specific resting state networks to respiration (Figs. 3 and 4), suggesting that these amplitude correlations within resting state networks are at least partially related to respiration.

### Active sensing, respiration, and behaviour

The widespread extent of the RMBO network critically corroborates previous suggestions of respiration as an overarching ‘clock’ mechanism organising neural excitability throughout the brain ^13^. Excitability adapts neural responses to current behavioural demands, which is why respiratory adaptation to such demands in animals ^39^ and humans ^40^ have accordingly been interpreted as functional body-brain coupling: With cortical excitability fluctuating over the respiration cycle, information sampling and motor execution during phases of high excitability optimises efficient communication between brain areas and/or effector muscles. Indeed, animals as well as humans appear to actively align their breathing to time points of particular behavioural relevance for the sake of efficiency through optimised neural processing. Consequently, human respiration has fittingly been cast as *active sensing* ^41^, adopting key premises from predictive brain processing accounts ^42^ to explain how respiration synchronises time frames of increased cortical excitability with the sampling of sensory information. Our results provide first insights into how established mechanisms like crossfrequency phase-amplitude coupling (in this case, coupling peripheral to neural rhythms) are implemented on a global scale to translate respiratory rhythms into neural oscillations of various frequencies and how the resulting anatomical pattern of RMBOs reflects spectral specificity.

### Potential mechanisms behind RMBOs

Cross-frequency coupling is widely regarded as the core mechanism of translating slow rhythms into faster oscillations and has conclusively been shown to be driven by respiratory rhythms within the OB in mice ^13^. Here, slow respiration-induced bulbar rhythms are transmitted through piriform cortex and subsequent cortico-limbic circuits to modulate the amplitude of faster oscillations in upstream cortical areas ^43^. With reference to the concept of active sensing introduced above, we argue that a similar case can be made for the cerebellum: There is broad consensus that the cerebellum is involved in computations attributed to *internal forward models*, predominantly in motor control ^44^. These forward models refer to an internal representation of potential outcomes of an action in order to compare estimated and actual consequences. Importantly, they are just as crucial for perception and cognition as they are for motor performance, leading to the suggestion that cerebellar processing may help to align and adaptively modify cognitive representations for skilled and error-free cognitive performance ^34^. One way in which respiration potentially modulates global computations within the RMBO network is through cross-frequency coupling, meaning that slow respiratory rhythms drive slow neural oscillations which are subsequently translated into faster cortical rhythms.

Strong support for this hypothesis comes from a recent study ^45^ showing that the cortical readiness potential, originating within premotor areas, fluctuates with respiration. Notably, the authors suggest cross-frequency coupling to involve neural interactions between premotor areas and both insular and cingulate cortex as well as the medulla, which is precisely the pathway we propose to connect deep and cortical nodes within the RMBO network. A simple graph model of excitatory and inhibitory cells has been shown as proof of principle for cortical gamma modulation through respiration (modelled as sinusoidal input) ^46^. The authors later concluded that respiration-locked brain activity has two driving sources ^47^: On the one hand, respiration entrains OB activity via mechanoreceptors (i.e., phase-phase coupling), as seen in LFP ^25^. On the other hand, Heck and colleagues propose extrabulbar sources within the brain stem. As argued above, the preBötzinger complex and connected medullar nuclei constitute an ideal candidate to drive high-frequency oscillations through respiratory rhythms (i.e., phase-amplitude coupling ^13^). Functionally, respiration thus appears to modulate higher oscillatory frequencies (e.g. gamma) for the purpose of integrating locally generated assemblies across the brain ^48^. Our data now show that respiration-brain coupling i) spans an even more extensive network including deep cerebello-thalamic pathways and ii) involves a wider variety of oscillatory modulation than previously assumed. It remains an intriguing question as to which lower-level mechanisms are employed to drive respiration-dependent changes in oscillatory network activity.

One potential physiological mechanism depends on arterial CO_2_ levels, usually operationalised as the CO_2_ gas partial pressure at the end of exhalation (p_ET_CO_2_). Natural fluctuations in arterial CO_2_ during normal breathing were shown to be sufficient to significantly influence neural oscillatory power in delta, alpha, beta, and gamma frequencies ^49^. Most likely, this dependence is of neurochemical origin through an inverse relationship between the concentration of arterial CO_2_ and pH ^50^. In turn, pH is mediated by extracellular adenosine, whose release acts to reduce cortical excitability during hypercapnia ^51^. This explanation fits well with previous reports showing that inspiration increases cortical excitability (see above), a link that appears to be exploited in self-paced protocols (e.g., spontaneous inspiration at behavioural responses ^7^) and has proven beneficial for performance, e.g. in cognitive processing ^9^ and motor control ^10^.

While the RMBO network presented here provides the most comprehensive account of human respiration-brain coupling to date, central research questions emerge as objectives for future work. First, having established the sources of respiration-related changes to neural oscillations, the transition from resting state to task context will illuminate the relevance of RMBOs for behaviour. Cognitive, perceptive, and motor performance have been shown to be modulated by respiration, warranting a closer assessment of the where (i.e., which site) and when (i.e., at which phase) of task-related RMBOs. Second, we have outlined functional pathways connecting the cerebellum to cerebral cortex via medullar and thalamic projections as well as the close link between olfactory bulb and parahippocampal as well as prefrontal cortices. These putative hierarchies should be tested with directional measures of functional connectivity in order to reveal organisational relations within the RMBO network. Similarly, directed connectivity analysis can disambiguate bottom-up from topdown signals within the wider RMBO network and potentially illuminate the notably lateralised effects within it: Although lateralisation is not uncommon in well-established functional networks (e.g. related to attention; see ^52^, it will be instructive for future work to validate whether RMBOs reliably prove stronger in one hemisphere (as was the case for insular and FEF) or whether there is a separate dynamic underlying the involvement of individual nodes.

In summary, our comprehensive investigation of respiration-brain coupling emphasizes respiration as a powerful predictor for amplitude modulations of rhythmic brain activity across diverse brain networks. These modulations are mediated by cross-frequency coupling (linking respiratory to neural rhythms) and encompass all major frequency bands that are thought to differentially support cognitive brain functions. Furthermore, respiration-brain coupling extends beyond the core respiratory control network to well-known resting-state networks such as default mode and attention networks. Our findings therefore identify respiration-brain coupling as a pervasive phenomenon and underline the fact that body and brain functions are intimately linked and, together, shape cognition.

## Acknowledgements

This work was supported by the Interdisciplinary Center for Clinical Research (IZKF) of the medical faculty of Münster (Gro3/001/19). JG was further supported by the DFG (GR 2024/5-1).

## Methods

### Participants

Twenty-eight volunteers (14 female, age 24.8 ± 2.87 years [mean ± SD]) participated in the study. All participants denied having any respiratory or neurological disease and gave written informed consent prior to all experimental procedures. The study was approved by the local ethics committee of the University of Muenster and complied with the Declaration of Helsinki.

### Procedure

Participants were seated upright in a magnetically shielded room while we simultaneously recorded respiration and MEG data. MEG data were acquired using a 275 channel whole-head system (OMEGA 275, VSM Medtech Ltd., Vancouver, Canada) and continuously recorded at a sampling frequency of 600 Hz. During recording, participants were to keep their eyes on a fixation cross centred on a projector screen placed in front of them. To minimise head movement, participants’ heads were stabilised with cotton pads inside the MEG helmet. Data were acquired in two runs of 5 min duration with an intermediate self-paced break. Participants were to breathe automatically while tidal volume was measured as thoracic circumference by means of a respiration belt transducer (BIOPAC Systems, Inc., Goleta, United States) placed around their chest.

For MEG source localisation we obtained high-resolution structural MRI scans in a 3T Magnetom Prisma scanner (Siemens, Erlangen, Germany). Anatomical images were acquired using a standard Siemens 3D T1-weighted whole brain MPRAGE imaging sequence (1 x 1 x 1 mm voxel size, TR = 2130 ms, TE = 3.51 ms, 256 x 256 mm field of view, 192 sagittal slices). MRI measurement was conducted in supine position to reduce head movements and gadolinium markers were placed at the nasion as well as left and right distal outer ear canal positions for landmark-based co-registration of MEG and MRI coordinate systems. Data preprocessing was performed using Fieldtrip ^53^ running in Matlab R2018b (The Mathworks, Inc., Natick, United States). Both MEG and respiration data were resampled to 300 Hz prior to further analyses.

### MRI co-registration

Co-registration of structural MRIs to the MEG coordinate system was done individually by initial identification of three anatomical landmarks (nasion, left and right pre-auricular points) in the participant’s MRI. Using the implemented segmentation algorithms in Fieldtrip and SPM12, individual head models were constructed from anatomical MRIs. A solution of the forward model was computed using the realistically shaped single-shell volume conductor model^54^ with a 5 mm grid defined in the MNI template brain (Montreal Neurological Institute, Montreal, Canada) after linear transformation to the individual MRI.

### Computation of global field power

For the computation of global field power, the time courses of each channel of each participant were individually subjected to a continuous wavelet transform using a Morlet wavelet for 36 frequencies (from 2 Hz to 20 Hz in steps of 2 Hz and then in steps of 5 Hz up to 150 Hz). Next, we computed the absolute value of this complex-valued data and averaged these amplitude values across channels.

### Head movement correction

In order to rule out head movement as a potential confound in our analyses, we used a correction method established by Stolk and colleagues ^55^. This method uses the accurate online head movement tracking that is performed by our acquisition system during MEG recordings. This leads to six continuous signals (temporally aligned to the MEG signal) that represent the x, y, and z coordinates of the head center (H_x_, H_y_, H_z_) and the three rotation angles (H_φ_, H_θ_, H_ψ_) that together fully describe the head movement. Head movement correction was performed by computing a general linear model (GLM)

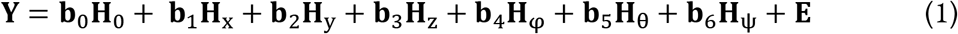

estimating the coefficients b and computing the residual signal Y. This effectively removes signal components that can be explained by translation or rotation of the head with respect to the sensors. We used this correction on each individual MEG sensor signal (and individual voxel signals for source-space analyses) as part of the preprocessing.

### Computation of modulation index and phase-triggered average

The modulation index (MI) quantifies cross-frequency coupling and specifically phase-amplitude coupling ^24^. Here, it was used to study to what extent the amplitude of brain oscillations at different frequencies is modulated by the phase of respiration. To this end, the instantaneous phase of the respiration time course was computed with the hilbert transform. Next, the time series at each sensor location were sequentially subjected to a continuous Morlet wavelet transformation at frequencies ranging from 2-150 Hz (with 2 Hz spacing below 20 Hz and 5 Hz spacing above 20 Hz) using the *cwtft* function in Matlab with default settings. This function computes a continuous Morlet wavelet transform using a Fourier transform-based algorithm. The Fourier transform of our wavelet is defined as

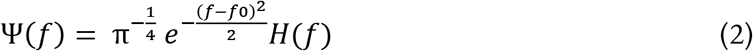

where H(f) is the Heaviside function and f0 is the center frequency in radians/sample. We then computed the amplitude envelope and smoothed it with a 300ms moving average. MI computation was then based on the average amplitude at 20 different phases of the respiratory cycle. Any significant modulation (i.e. deviation from a uniform distribution) is quantified by the entropy of this distribution. To account for frequency-dependant biases we followed previously validated approaches ^9,56^ and computed 200 surrogate MIs using random shifts of respiratory phase time series with concatenation across the edges. The normalised MI was computed by subtracting, for each frequency, the mean of all surrogate MIs and dividing by their standard deviation leading to MI values in units of standard deviation of the surrogate distribution (see Fig. 1b). Visual inspection confirmed that this removed the frequency bias in raw MI values (stronger MI for low frequencies compared to high frequencies). The computation resulted in normalised MI values for each voxel, frequency, and participant. Following the established approach by Maris and colleagues ^57^, significance of these normalised MI values on the group level was determined by means of cluster-based permutation testing. Specifically, we conducted a series of one-tailed t-tests of the group-level average MI at each frequency against the 95th percentile of the null distribution constructed from a Monte Carlo approximation. For this approximation, we constructed random permutations of single-subject MI x frequency matrices and computed the maximum of the cluster-level summed t-values. After repeating the randomisation procedure 5000 times, the original group-level MI values (per voxel and frequency) were compared to the histogram of the randomised surrogate data. Clusters in the original data were deemed significant when they yielded a larger test statistic than 95% of the randomised surrogate data. Cluster-level results were subsequently FDR-corrected across frequencies at a significance level of α = .05.

To assess oscillatory modulation over time, the phase-triggered average (PTA) was computed from the smoothed, band-specific amplitude envelopes averaged across all sensors. Time points of peak inhalation were detected from the respiration phase angle time series using Matlab’s *findpeaks* function. For each time point of peak inhalation, global field power across all 36 frequencies was averaged within a time window of ± 1000 samples centered around peak inhalation. The resulting 36 frequencies x 2000 samples matrix was finally normalised across the time dimension, leading to z-scores of whole-head oscillatory power phase-locked to the respiration signal. This analysis is equivalent to a wavelet-based time-frequency analysis. Computations were done separately for both MEG runs, normalised across the time domain, and finally averaged across runs and participants.

### Extraction of time series in source space

Source reconstruction was performed using the linearly constrained minimum variance (LCMV) beamformer approach ^58^, where the lambda regularisation parameter was set to 0%. This approach estimates a spatial filter for each location of the 5-mm grid along the direction yielding maximum power. The sensor covariance matrix used for the LCMV-beamformer was computed across the whole data set.

### Rank optimisation and non-negative matrix factorisation

In our efforts to anatomically localise respiration phase-dependent modulation effects, we employed a spatially sparse variant of non-negative matrix factorisation to reduce the high (voxelwise) dimensionality in our data. Sparse NMF allows us to describe modulation indices across the brain as a low-dimensional combination of locally constrained network components, each of which provides a spectral profile for each participant. As the modulation index is inherently non-negative and the interpretation of NMF output matrices is rather straightforward, non-negative factorisation in general was well-suited to meet our demands. The sparse factorisation approach in particular has two key advantages over a regular NMF approach: First, regarding network identification, the sparsity constraints are highly beneficial in obtaining spatially specific rather than broad topologies, which was central for the next steps of our analyses. Second, these spatially specific topologies greatly enhance the precision with which time x frequency modulation characteristics can be displayed within one network component – the more distant voxels are included, the more component-specific modulations are diluted. In order to balance baseline differences between participants in preparation of the NMF, modulation index matrices of all 28 participants (20,173 voxels x 36 frequencies) were first normalised by their standard deviation ^59^. These matrices were then averaged across both runs to yield one average matrix per participant. Individual matrices were transposed and concatenated to form one group-level input matrix (1008 [frequencies x participants] x 20,137 voxels) for the NMF. To determine the number of main components to be extracted from NMF, we used the *choosingR* Matlab function ^60^ that chooses the optimal rank based on singular value decomposition. Specifically, the function extracts the singular values of a data matrix (in our case, participants’ MI matrices; size 36 frequencies x 20,173 voxels) and computes the sum of all non-zero elements of its diagonal. The optimal rank is then determined as the number of singular values that accounts for 90% of all diagonal entries. Applying this procedure to participants’ individual MI matrices (36 frequencies x 20,173 voxels) returned a dimensionality of 17 as the optimal desired number of network components for the subsequent NMF analysis.

Subsequently, we initialised the *sparsenmfnnls* algorithm from the NMF toolbox for Matlab ^61^. The algorithm factorises the concatenated input matrix **X** as

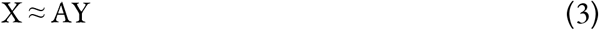

with the non-negative matrices A and Y aiming to minimise the following quantity:

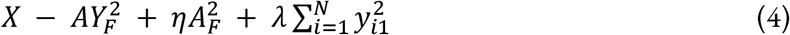

where *η* and *λ* are sparsity parameters. As NMF solutions vary as a function of their random starting position, we repeated this procedure 100 times and selected the best sparse solution based on its residuals. Two matrices were generated as the output of this procedure: First, the *basis matrix* **A** (1008 [frequencies x participants] x 17 components) represents the participant-specific spectral profile, effectively quantifying each participant’s relative contribution to the network components separately for each frequency. The basis matrix was reshaped to a 36 x 28 x 17 (frequencies x participants x network components) matrix for all further analyses. As the second NMF output, the *coefficient matrix* **Y** (17 components x 20,173 voxels) represents the spatial profile of the network components, quantifying each voxel’s relative contribution to the components.

### Component-level statistical analyses

While most components represented a single focal location due to the sparsity constraints embedded in the NMF algorithm, four components comprised distinct sub-networks of two or three anatomical sites. Spatial maps of all 17 network components were thresholded at *p* = .01 (see Fig. 2) and the resulting maps (volume ~200 voxels) were used as binary masks so that all further analyses were restricted to above-threshold voxels. To determine the frequency range(s) for which the modulation index within a particular component was significant on the group level, we once again followed a cluster-based permutation approach. To this end, we constructed a null distribution of MI using a Monte Carlo approximation with 5000 randomisation permutations. For each component, we conducted a series of one-tailed t-tests comparing the mean normalised MI at each frequency (averaged within the ROI) to the 95th percentile of the null distribution. The results were subsequently FDR-corrected for multiple comparisons at a significance level of α = .05 across all 36 frequencies and 17 components.

### Linear mixed effect models

We employed linear mixed effect modelling (LMEM) to investigate the relationship between the spatial organisation and spectral characteristics within the network of modulated components. LMEM models a response variable (in our case, modulation within a particular frequency band) as a linear combination of *fixed effects* shared across the population (i.e. anatomical coordinates of network components) and participant-specific *random effects* (i.e modulatory variation between participants). To assess potential links between spatial and spectral component properties, we first computed each component’s anatomical distance to the head centre as the vector norm of MNI coordinates in the x, y, and z plane:

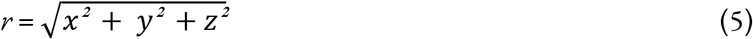

We specified two LMEMs to predict high frequency modulation indices (low gamma, high gamma) within each component as a function of its distance to the head centre, respectively:

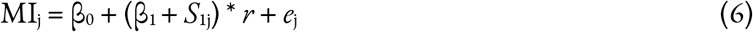

For participant *j*, the modulation index is expressed as a combination of the intercept (β_0_), the fixed effect of the component’s distance to the head centre (β_1_), and an error term (*e*_j_ ~ *N*(0,σ^2^)). We accounted for between-participant variation by specifying a random slope (*S*_1j_). An analogous approach was used to predict low frequency modulation indices (delta, theta) within each component as a function of its height in the sagittal plane (i.e. the z value of its MNI coordinates):

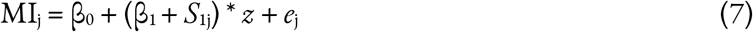

### Hierarchical clustering

Having localised the sources of global field power modulations within a constrained subset of anatomical sites, our next aim was to characterise these sources in terms of their spectro-temporal fingerprints. This way, we hoped to reveal systematic patterns of phase-locked oscillatory modulations over time and/or frequencies within the cortical and subcortical network. To this end, we first computed the group-level average matrix of modulation indices for 20173 voxels x 36 frequencies x 20 time bins. We used the anatomical distribution of each network component (thresholded at *p* = .01) to reduce this matrix to a component-specific ROI and aggregated 36 single frequencies into frequency bands as follows: delta (2-4 Hz), theta (4 – 8 Hz), alpha (8-12 Hz), beta (12-30 Hz), low gamma (30-70 Hz), and high gamma (70-150 Hz). This yielded one matrix (6 frequency bands x 20 time bins) per network component, all of which were concatenated to construct a distance matrix for the hierarchical clustering using the *hcluster* function within the Icasso toolbox for Matlab ^62^. This data-driven approach was employed to detect similarities of and differences between network components with regard to how oscillatory activity was modulated over the course of a respiration cycle. Following the suggested approach ^62^, we used visual inspection of the dendrogram and Matlab’s *evalclusters* function (with the *linkage* algorithm) to evaluate the clustering solutions. Based on a local maximum of the resulting silhouette value distribution, we settled on a total of seven clusters (see Fig. 3). We computed the average course of modulation indices over frequency bands within each cluster based on z-transformed spectral profiles of the contributing network components (as described above).

### Data availability

The anonymised data supporting the findings of this study are openly available from on the Open Science Framework (https://osf.io/6zbdu/).

