## Supplementary Information for "Respiration modulates oscillatory neural network activity at rest"

### Supplementary Information: Control analyses

#### Head movements

We conducted two types of control analyses with respect to head movements. First, based on previous work [11], it was reasonable to assume that there would be respiration-induced changes in head position and/or rotation. Therefore, we computed individual Spearman correlations between the normalised respiration time course and head movement traces of translation and rotation (in x, y, and z direction, respectively). Correlation coefficients were Fisher z-transformed and averaged across runs (for each participant) and across participants to yield group-level average correlation coefficients for all 6 head movement time courses. A series of t-tests revealed significant correlations between the respiration signal and translation in the x plane ($\rho(27) = -.16$, $t(27) = -10.31$, $p < .001$) as well as rotation in both x plane ($\rho(27) = -.16$, $t(27) = -10.99$, $p < .001$) and z plane ($\rho(27) = .16$, $t(27) = 11.08$, $p < .001$; all p-values corrected for multiple comparisons using the Bonferroni-Holm method). Supplementary Fig. 7 shows head movement traces (translation, rotation) phase-locked to respiration.

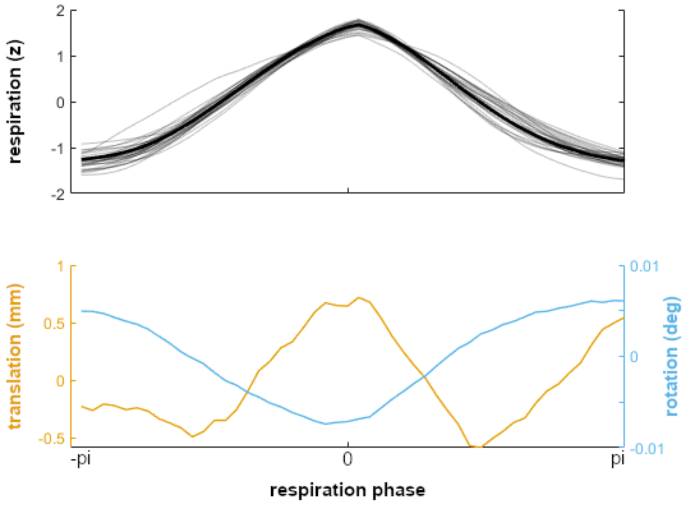


**Supplementary Fig. 7 | Head movement across the respiratory cycle.** Top panel shows individual (grey lines) and group-level average time courses of the normalised respiration signal (bold). Bottom panel shows group-level average head movement signals phase-locked to the respiration signal. Both translation (measured as Euclidean distance, yellow) and rotation (blue) are depicted as vector norms combining movement traces in x, y, and z directions.

As some correlation between respiration and head movement was to be expected, it was critical to use the regression approach described in the Methods section (see p. 26) to rule out that our results were confounded by head movements. Our second control analysis was conducted to make sure our head movement GLM did not miss any incremental influence of higher order head movement regressors on the MEG signal. To this end, we amended the original regression model to also contain non-linear regressors (up to the third order) and their derivatives in addition to the ‘raw’ time courses of translation and rotation. Slow drifts were removed by a third-order polynomial fit. We then repeated our sensor-level MI and PTA analyses with the expanded head movement correction and compared their results to those based on our original GLM (as reported in the Results section, see p. 5 and Fig. 1). We found the results to be virtually unchanged, indicating that our original head movement correction had picked up on all relevant portions of the signal:

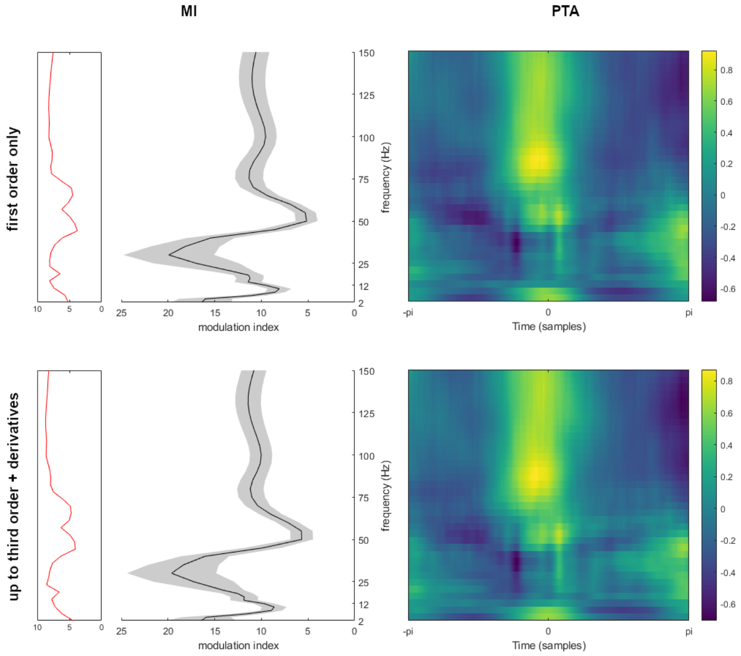


**Supplementary Fig. 8 | Comparison of sensor-level MI and PTA results for different head movement regression models.** The overall pattern MI and PTA results (upper panels, identical to Fig. 1b and c) was not altered by adding second and third order movement regressors plus their derivatives (lower panels).

Cluster permutation t-tests (with subsequent FDR correction for multiple comparisons, as implemented throughout the manuscript) confirmed that neither MI values (1 x 36 frequencies per participant) nor PTA values (36 frequencies x 2000 time samples per participant) showed any significant differences between the two regression models. Finally, to control for potential source-level effects, we subjected the total of 36 GLM regression weights for each participant (2 movements [translation, rotation] x 3 directions [x, y, z] x 3 orders [first, second, third] x 2 levels [raw, derivatives]) to a group-level analysis: Individual regression weights were averaged across both runs, yielding a matrix of 36 regression weights x 20,173 voxels per participant. Applying the same cluster permutation approach we followed throughout our analyses, we confirmed that there were indeed no clusters of voxels whose head movement regressors were significantly different from zero.

**High-frequency muscle artefacts**

Given the reported modulatory effects in the gamma band, it is crucial to rule out that they were caused by artificial, high-frequency muscle (EMG) activity. To this end, we conducted an ICA to isolate individual EMG components and repeated the analyses of MI and PTA on the time courses of these components. This analysis requires at least one ICA component reliably representing high-frequency EMG activity from most (if not all) participants in order to allow the estimation of group-level effects. Using Fieldtrip's 'ft_componentanalysis' and 'ft_icabrowser' functions (see www.fieldtriptoolbox.org for documentation), subject-level ICAs did not reveal a single component (within the 30 strongest components extracted in each run) whose topography, temporal distribution, and/or spectral characteristics suggested artificial EMG activity (following established criteria published in e.g. Muthukumaraswamy, 2013; Vigário et al., 2000; Mantini et al., 2008). In order to check for EMG artefacts in less powerful components, we re-ran the ICA to extract 60 components. Again, we failed to identify clear EMG components in the MEG signal.

The lack of EMG artefacts in our data is not surprising, given that a) participants were explicitly instructed to sit relaxed and refrain from clenching their teeth or neck, blinking etc and b) the task-free measurement made any and all muscle movements unnecessary. Furthermore, the manuscript cites a rather extensive body of literature demonstrating the link between respiration and gamma oscillations by means of phase-amplitude coupling (see the Discussion section, p. 14 ff).
