## Supplementary Figures for "Respiration modulates oscillatory neural network activity at rest"

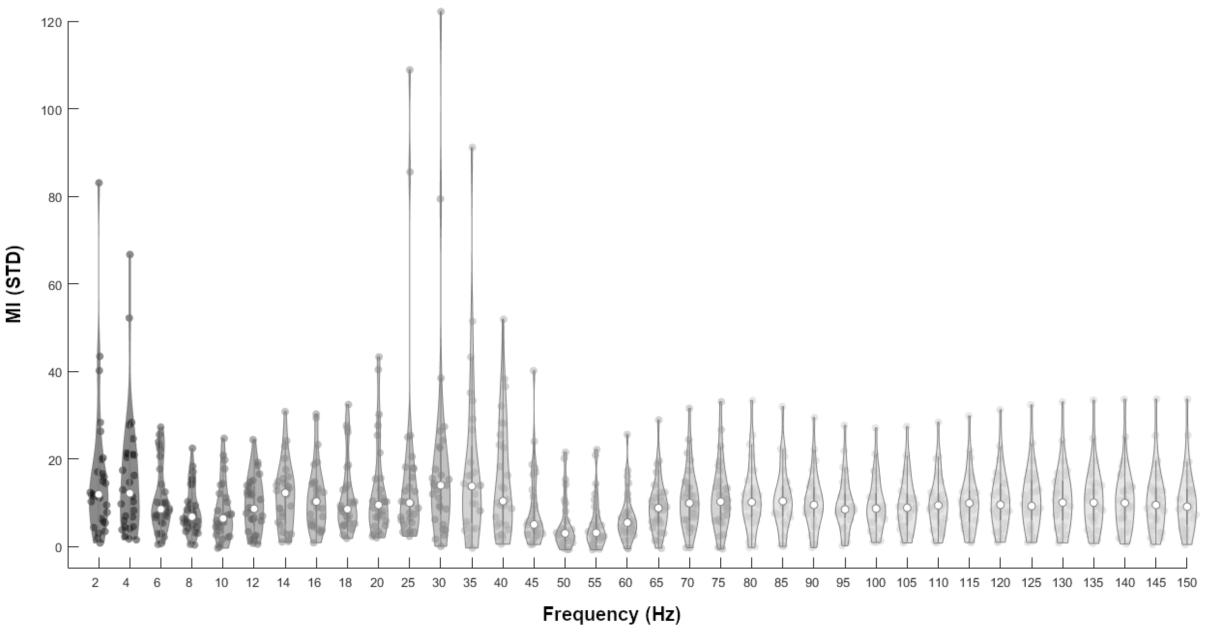


**Supplementary Fig. 1 | Range and distribution of sensor-level MI values.** Violin plot shows the distribution of individual MI values (depicted as dots) as well as the group-level median (white dot) across the whole frequency spectrum.

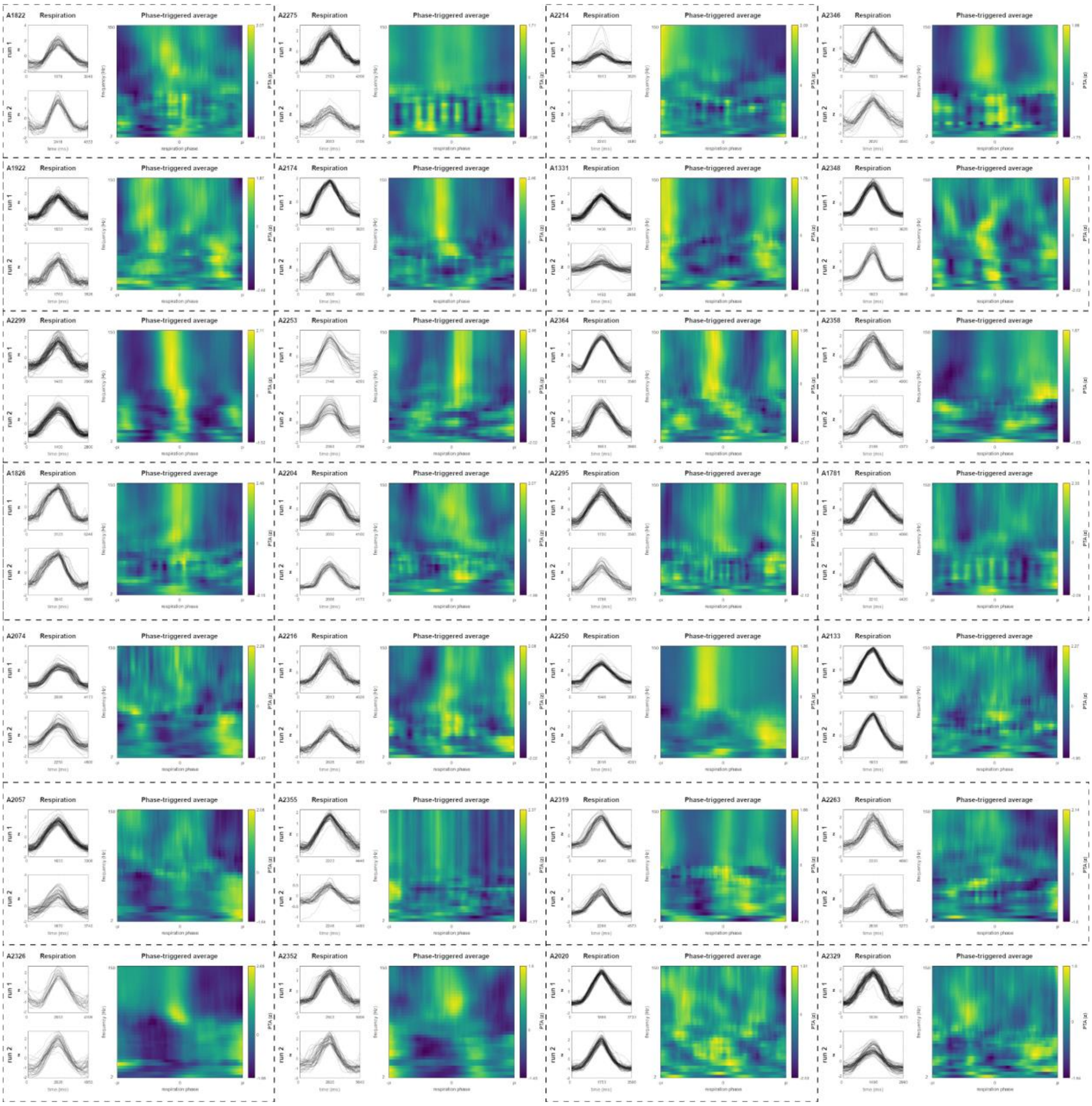


**Supplementary Fig. 2 | Individual respiration traces and phase-amplitude spectrograms.** Left panels show single respiration traces centred around peak inspiration from each run. Right panel shows the individual phase-amplitude spectrogram (averaged across all sensors). PTA values are shown as z-values, i.e. normalised within each frequency to reveal phase-related modulations.

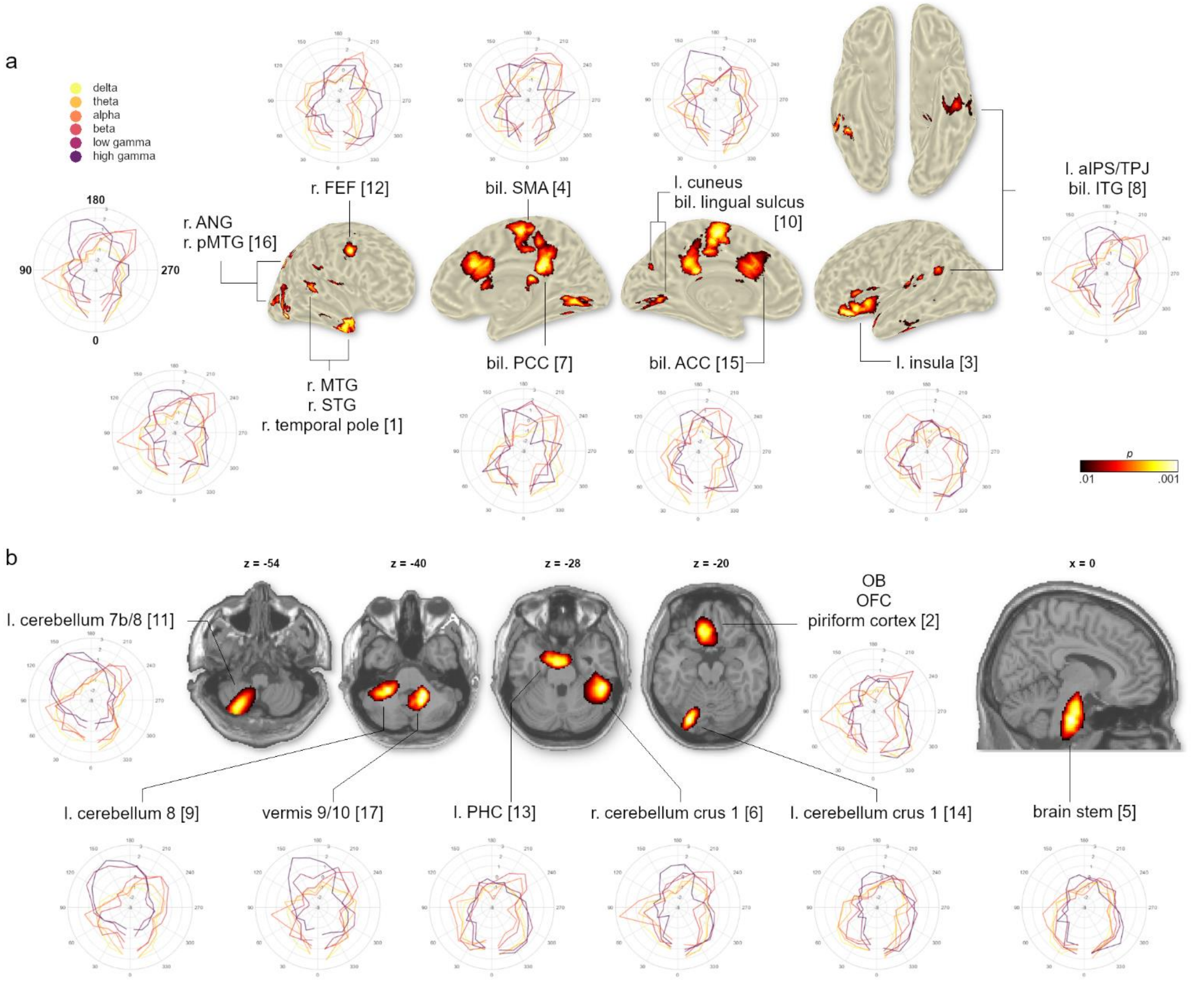


**Supplementary Fig. 3 | Temporal modulation profiles of NMF components whose neural oscillations were significantly modulated by respiration. a**, Cortical components plotted on an inflated brain surface. Polar plots show group-level normalized MI time courses averaged within frequency bands (delta to high gamma) over the entire respiration cycle. **b**, Subcortical components plotted on transverse and sagittal slices of the MNI brain. Same format as **a**.

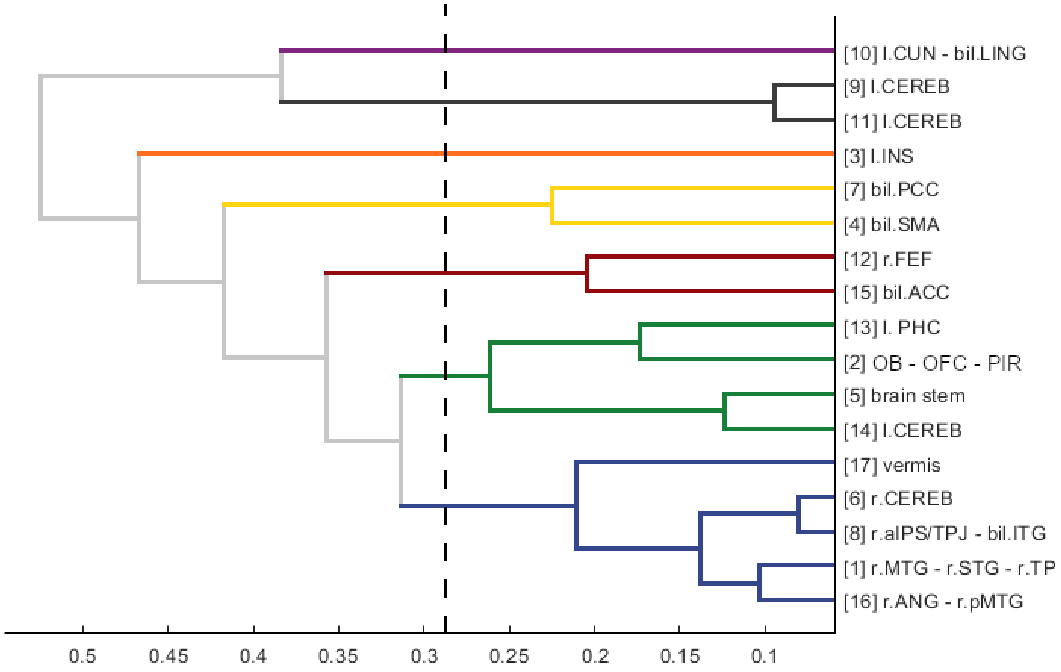


**Supplementary Fig. 4 |** Dendrogram of the hierarchical clustering performed on all 17 main components from the NMF analysis. Dashed vertical line illustrates the cut-off criterion, yielding a total of 7 clusters. Cluster colouring is identical to Figures 3 and 4.

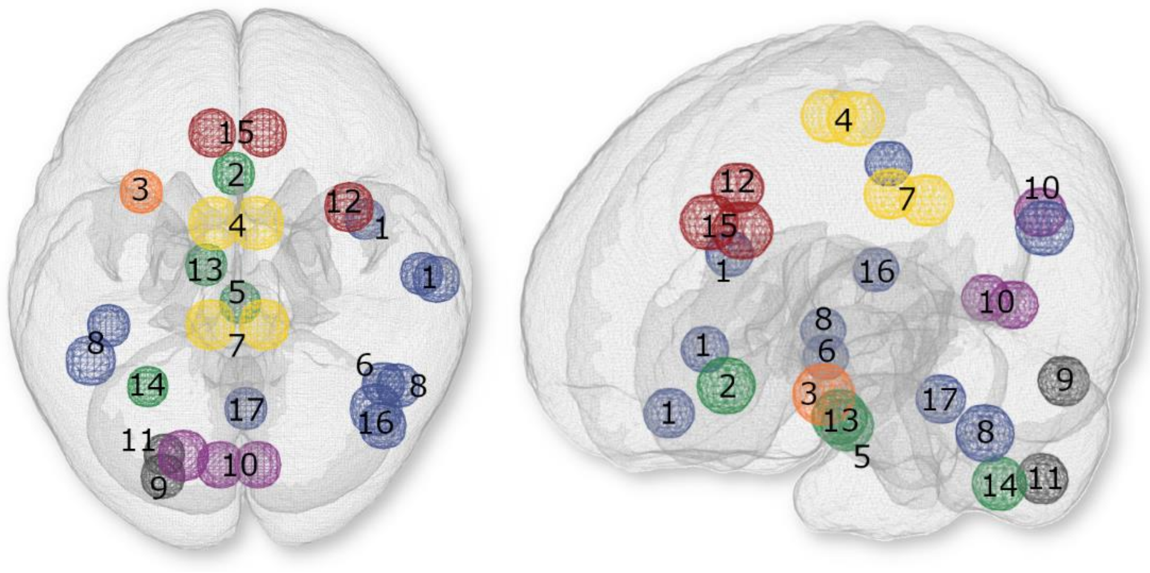


**Supplementary Fig. 5 |** Top (left) and side view (right) of the RMBO network spanned by the 17 significant NMF components. Numbering corresponds to Figures 2 and 4c as well as Supplementary Fig. 1. Cluster colouring is identical to Figures 3 and 4 as well as Supplementary Fig. 4.

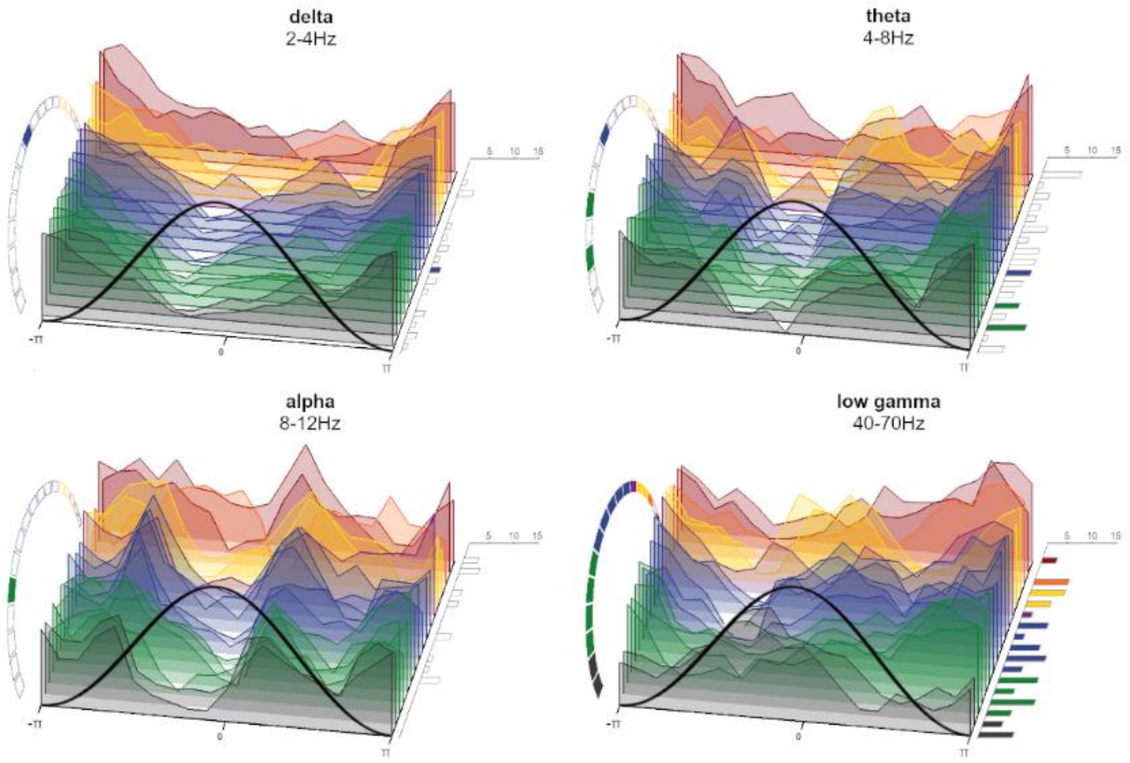


**Supplementary Fig. 6 |** Waterfall plots show z-transformed amplitude modulation phase-locked to the respiration cycle for the remaining frequency bands not shown in Fig. 4d. Clusters of NMF components are shown in the same order as in Fig. 4c. Right-panel bar graphs show the number of participants whose modulation within the respective component was strongest for the depicted frequency band (vs all other frequency bands). Coloured bars and circular segments mark NMF components for which the respective frequency band was significantly modulated by respiration phase.
